# Comparative proteomics uncovers low asparagine insertion in *Plasmodium* tRip-KO proteins

**DOI:** 10.1101/2023.08.09.552632

**Authors:** Martina Pitolli, Marta Cela, Delphine Kapps, Johana Chicher, Laurence Despons, Magali Frugier

## Abstract

tRNAs are not only essential for decoding the genetic code, but their abundance also has a strong impact on the rate of protein production, folding, and on the stability of the translated messenger RNAs. *Plasmodium* expresses a unique surface protein called tRip, involved in the import of exogenous tRNAs into the parasite. Comparative proteomic analysis of the blood stage of wild-type and tRip-KO variant of *P. berghei* parasites revealed that down-regulated proteins in the mutant parasite are distinguished by a bias in their asparagine content. Furthermore, the demonstration of the possibility of charging host tRNAs with *Plasmodium* aminoacyl-tRNA synthetases, led us to propose that, imported host tRNAs participate in parasite protein synthesis. These results also suggest a novel mechanism of translational control in which import of host tRNAs emerge as regulators of gene expression in the *Plasmodium* developmental cycle and pathogenesis, by enabling the synthesis of asparagine-rich regulatory proteins that efficiently and selectively control the parasite infectivity.

## Introduction

We have discovered the only example to date of an exogenous tRNA import pathway, summarized in Fig. 1 ^1–4^. In *Plasmodium*, the malaria parasite, *P. berghei* sporozoites (the extracellular infective form of the parasite, Supplementary Fig. S1a) isolated from the salivary glands of infected mosquitoes import exogenous tRNAs *via* a surface protein, named tRip (tRNA import protein). However, the absence of homologous mechanisms in other organisms raises the question of the role of this unique tRNA import and its mode of action. It has been established that (i) tRip is a homodimeric protein made of an N-terminal GST-like domain and a C-terminal EMAPII-like tRNA binding domain; (ii) tRip binds human tRNAs with high affinity *in vitro* and with a stoichiometry of one tRNA per tRip dimer; (iii) tRip is anchored to the parasite’s plasma membrane with its tRNA binding domain exposed to the host system; (iv) immunolocalization experiments found tRip expressed both in the liver and blood stages in the vertebrate host, as well as in the intestine and salivary glands of the mosquito; (v) *in vitro*, exogenous tRNAs enter living sporozoites; (vi) the knockout parasite, tRip-KO, does not import tRNAs, its protein biosynthesis is significantly reduced, and its growth is decreased in vertebrate blood compared to the wild-type parasite.

**Figure 1.**
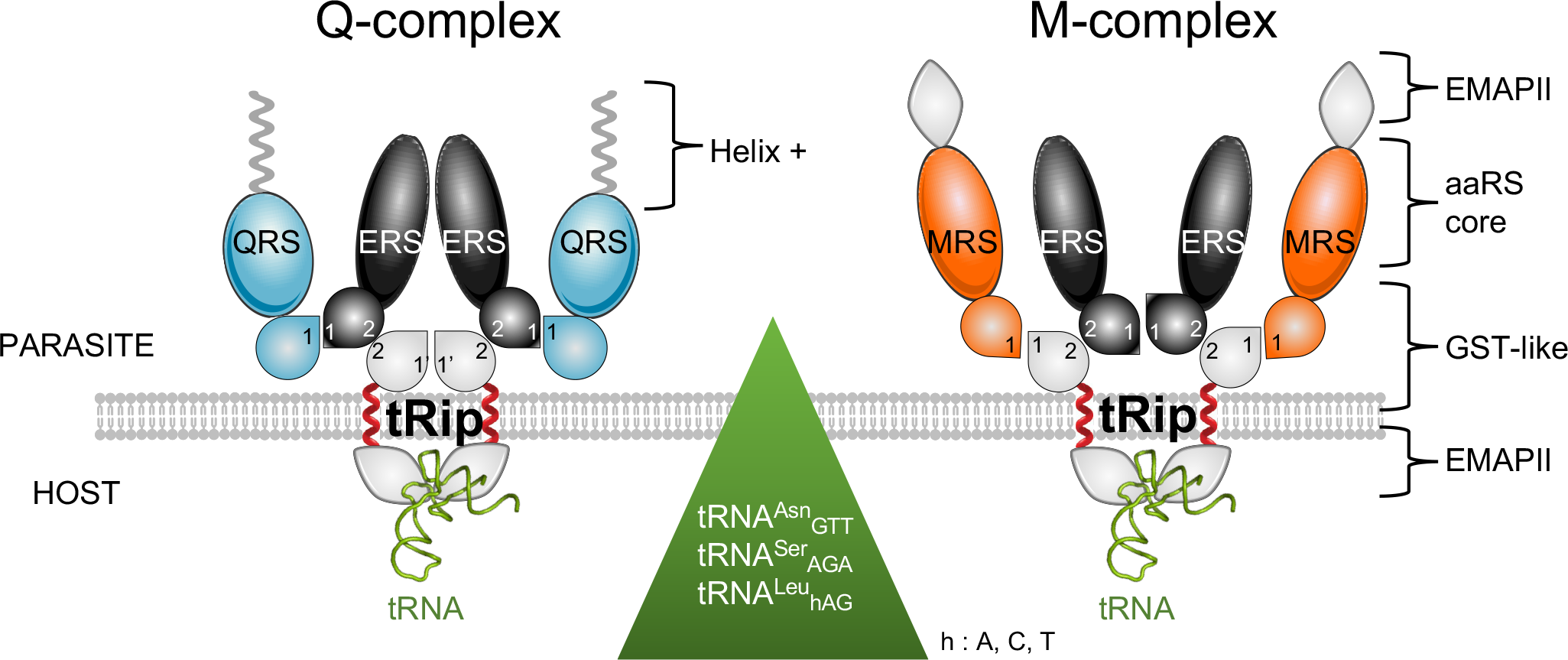
*P. berghei* membrane-bound multi-synthetase complexes. *Plasmodium* is characterized by the presence of two multisynthetase complexes (MSC) named Q-complex and M-complex. The Q-complex is composed of glutamyl- (ERS) and glutaminyl- (QRS) tRNA synthetases linked to a dimer of tRip whereas the M-complex is composed of tRip and Methionyl-tRNA synthetase (MRS) organized around a dimer of ERS. tRip is therefore an AIMP (Aminoacyl-tRNA synthetase Interacting Multifunctional Protein). tRip, ERS, QRS, and MRS are schematized and colored in grey, black, cyan, and orange, respectively. the GST-like domains are shown as a drop and the C-terminal domains of tRip, QRS and MRS involved in tRNA binding are either EMAPII-like domains (tRip and MRS, grey diamonds) or a positively charged α-helix (QRS, shown as a grey helix). The characteristic feature of *Plasmodium* MSCs is that tRip is a membrane protein (the transmembrane helix is shown in red) with the GST-like domain required for MSC formation localized inside the parasite and the tRNA binding domain exposed outside the parasite to host tRNAs. This unique organization justifies that only the EMAPII-like domain of tRip (tRip_200-402_) is fused at the C-terminal domain of a GST domain and used as a target for the selection of aptamers capable of inhibiting tRip tRNA binding. Interfaces 1, 1’ or 2, involved in protein-protein interactions, are indicated in the corresponding GST-like domains. For the sake of simplicity, this illustration does not show the parasitophorous membrane that surrounds the parasite in the infected red blood cell.

Recently, the crystal structure of the dimeric N-terminal GST-like domain of *P. vivax* tRip was solved and revealed a unique homodimerization interface ^5^. We confirmed by SAXS that this unusual interface exists in solution and that it allows Multi-Synthetase Complex (MSC) formation ^3,4^. Indeed, tRip is an aminoacyl-tRNA synthetase interacting multifunctional protein (AIMP) since it specifically co-immunoprecipitate with three aminoacyl-tRNA synthetases (aaRS), namely glutamyl- (ERS), glutaminyl- (QRS) and methionyl- (MRS) tRNA synthetases; These aaRSs all contain an N-terminal GST-like domain involved in MSC assembly. Unexpectedly, these proteins form two exclusive heterotrimeric MSCs with a 2:2:2 stoichiometry: a Q-complex (tRip:ERS:QRS) and an M-complex (tRip:ERS:MRS) characterized by different biophysical properties and interaction networks (Fig. 1). We could also identify a set of host tRNAs preferentially bound by tRip (Fig. 1) and potentially best imported into the parasite. Interestingly, tRip does not bind to tRNAs in a sequence-dependent manner, but would rather recognize post-transcriptional modifications that modulate this interaction ^2^. In contrast to what has been shown for its cytosolic homologue Arc1p in *Saccharomyces cerevisiae* ^6^, the tRNAs that best bind tRip do not match the aaRSs that compose MSCs. It suggests that, although tRip is an AIMP, it is not dedicated to the aminoacylation of specific tRNAs (2), thereby leading us to search for a novel function for this unique membrane protein and the tRNA import with which it is associated.

The AIMP tRip is not the only protein that has been characterized as cytosolic in other organisms and is localized on the surface of *Plasmodium*. This is also the case for other RNA-binding proteins such as the poly-A binding protein-1 (PABP-1) ^7^ and the glyceraldehyde-3 phosphate dehydrogenase (GAPDH) ^8^, which are both found on the surface of the sporozoites. The presence of different RNA-binding proteins at the parasite-host interface is another indication that RNA exchange might control unsuspected host-parasite interactions that take place during the parasite life cycle. In the present study, we compared the proteomes of the wild-type (WT) and the tRip-KO blood stage parasites with the objective of understanding the fate of imported tRNAs in *Plasmodium* protein synthesis and its infectious process.

## Results

### Host tRNAs are aminoacylated by the cognate parasite aaRSs *in vitro*

Cross-aminoacylation reactions were tested using the aspartyl-, tyrosyl-, and asparaginyl-tRNA synthetases from *H. sapiens* and *P. falciparum* (Supplementary Fig. S2c) *in vitro* to determine their ability to aminoacylate human tRNA^Asp^, tRNA^Tyr^, and tRNA^Asn^, respectively using crude human tRNA (Fig. 2). All three *Plasmodium* aaRSs can aminoacylate nearly the same level of tRNA as their human counterparts, although asparaginylation by the parasite enzyme is effective on about 80% of human tRNA^Asn^ isodecoders. It indicates that host tRNAs, when imported into parasites by tRip can be aminoacylated by parasite aaRSs to be used in protein translation.

**Figure 2.**
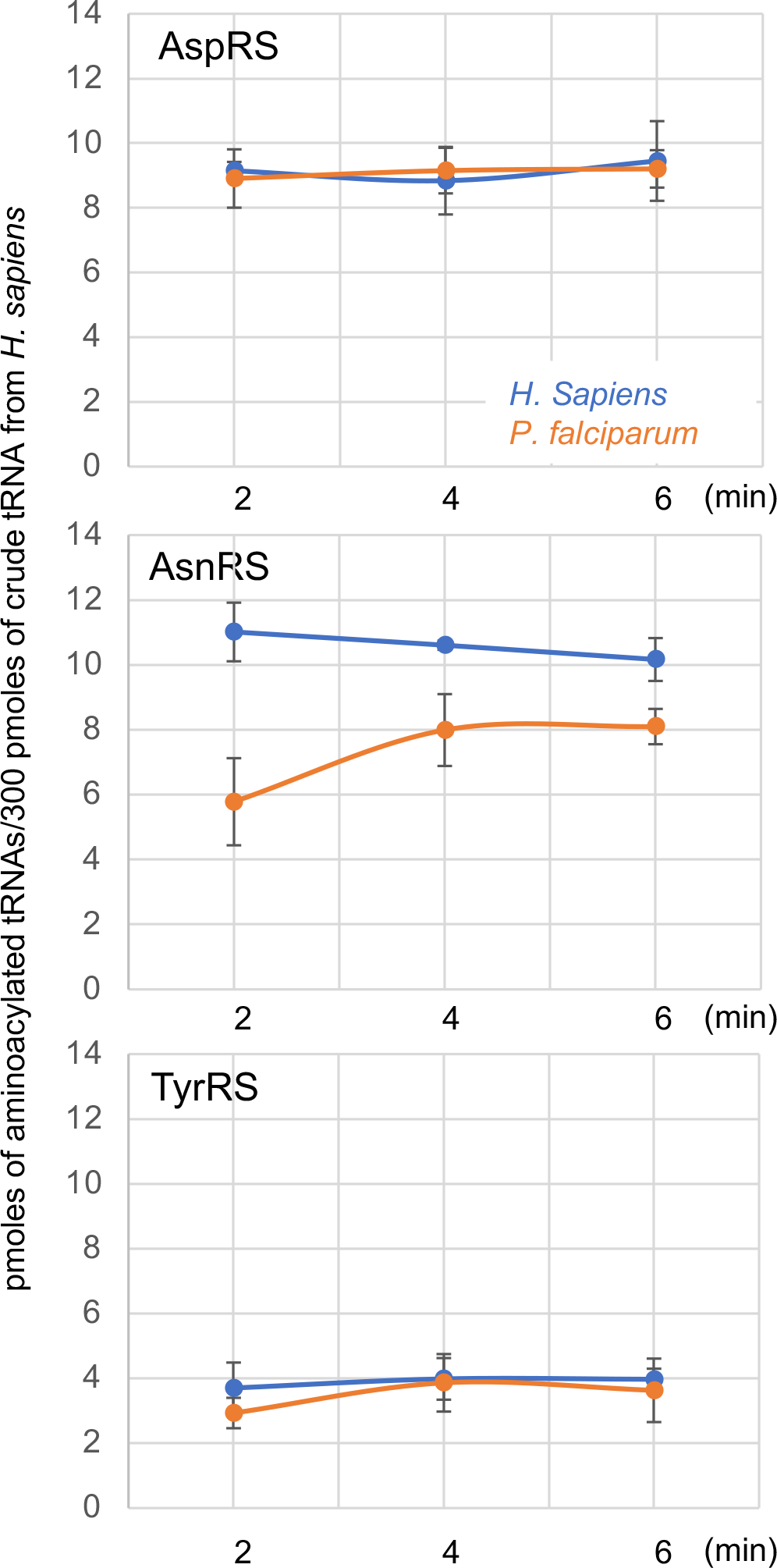
Detection of host tRNAs in WT parasites and cross aminoacylation of host tRNAs by the parasite aminoacyl-tRNA synthetases. (**a**) One µg of total RNA from the liver of non-infected mice and 1 µg of total RNA from WT and KO parasites were analyzed on an ethidium bromide-stained 12% denaturing analytical gel. L corresponds to a DNA ladder. Northern blot experiments were completed with either probes designed to detect *P. berghei* tRNAs (*Pb*-^Glu^_TTC_, *Pb*-^His^, *Pb*-^Met^_i_ and *Pb*-^Met^_e_) or *M. musculus* tRNAs (*Mm*-^Glu^_TTC_, *Mm*-^His^, *Mm*-^Met^ _i_ and *Mm*-^Met^ _e_) (Supplementary Fig. S2). The four tRNAs belong to the category of tRNAs that interact most strongly with tRip *in vitro*. Detection of *Mm* tRNAs by Western blots necessitated extensive washing and long exposure of the films. (**b**) Comparative aminoacylation plateaus. Six aaRSs, aspartyl-, asparaginyl- and tyrosyl-tRNA synthetases from *H. sapiens* (in blue) or *P. falciparum* (in orange) were tested with crude human tRNAs under the same experimental conditions (6 µM crude tRNA and 0.2 µM aaRSs). Aminoacylation plateaus were measured at 2, 4, 6 min of incubation. Human crude tRNAs are potentially transcribed from 420 genes, amongst which 19 genes encode tRNA^Asp^, 34 genes encoding tRNA^Asn^ and 15 genes encoding tRNA^Tyr 40^. Error bars represent the standard deviation (SEM) of three independent experiments.

### Comparative proteomics of tRip-KO *versus* wild-type (WT) schizonts

We investigated the relative protein abundance of tRip-KO *versus* WT in the schizont stage of synchronized parasites (Supplementary Fig. S1b) using liquid chromatography coupled to tandem mass spectrometry (LC-MS/MS) and a label-free quantification method. To highlight the most relevant effects associated with the deletion of tRip, we chose to compare equal amounts of proteins, even though tRip-KO parasites grow slower than wild-type parasites and their translation efficiency is significantly reduced ^1^. Three biological replicates for tRip-KO and WT were defined for the schizonts samples (Supplementary Fig. S4). The amounts of proteins (KO and WT) were comparable as most proteins were stable in the proteomic data. This is notably the case for four constitutively highly expressed proteins: EF1-α (PBANKA_1133300), Hsp70 (PBANKA_0914400), enolase (PBANKA_1214300) and histone H4 (PBANKA_941900) (Supplementary Fig. S4). Protein abundance was calculated using the intensity of *P. berghei* peptides, which identified between 347 and 469 proteins. Metric multidimensional scaling (MDS) (included in Supplementary Fig. S4) indicated clustering of the tRip-KO and WT parasites, suggesting there are distinct groups of differentially abundant proteins between the two genotypes. Among these proteins, only 8 were significantly up-regulated and 49 were down-regulated in the tRip-KO parasites compared to the WT (*p-value* < 0.05) with a at least a 2-fold change in abundance; they are highlighted in the volcano plots in orange and green, respectively (Fig. 3a). Most of down-regulated proteins are involved in translation (31% ribosomal proteins, initiation, and elongation factors); the two other abundant categories were surface proteins involved in transport and invasion (16%) and proteins with unknown functions (14%) (Supplementary Fig. S4). Interestingly, the deregulated proteins include several aminoacyl-tRNA synthetases. The glutamyl-tRNA synthetase (ERS), which is tightly associated to tRip in the two plasmodium MSCs is reduced in the tRip-KO sample and both the lysyl- (KRS) and the asparaginyl (NRS) -tRNA synthetases are increased in the mutant compared to the WT parasite (Supplementary Fig. S4), NRS and ERS being just below the threshold with *p-values* of 0.0723 and 0.0627, respectively.

**Figure 3.**
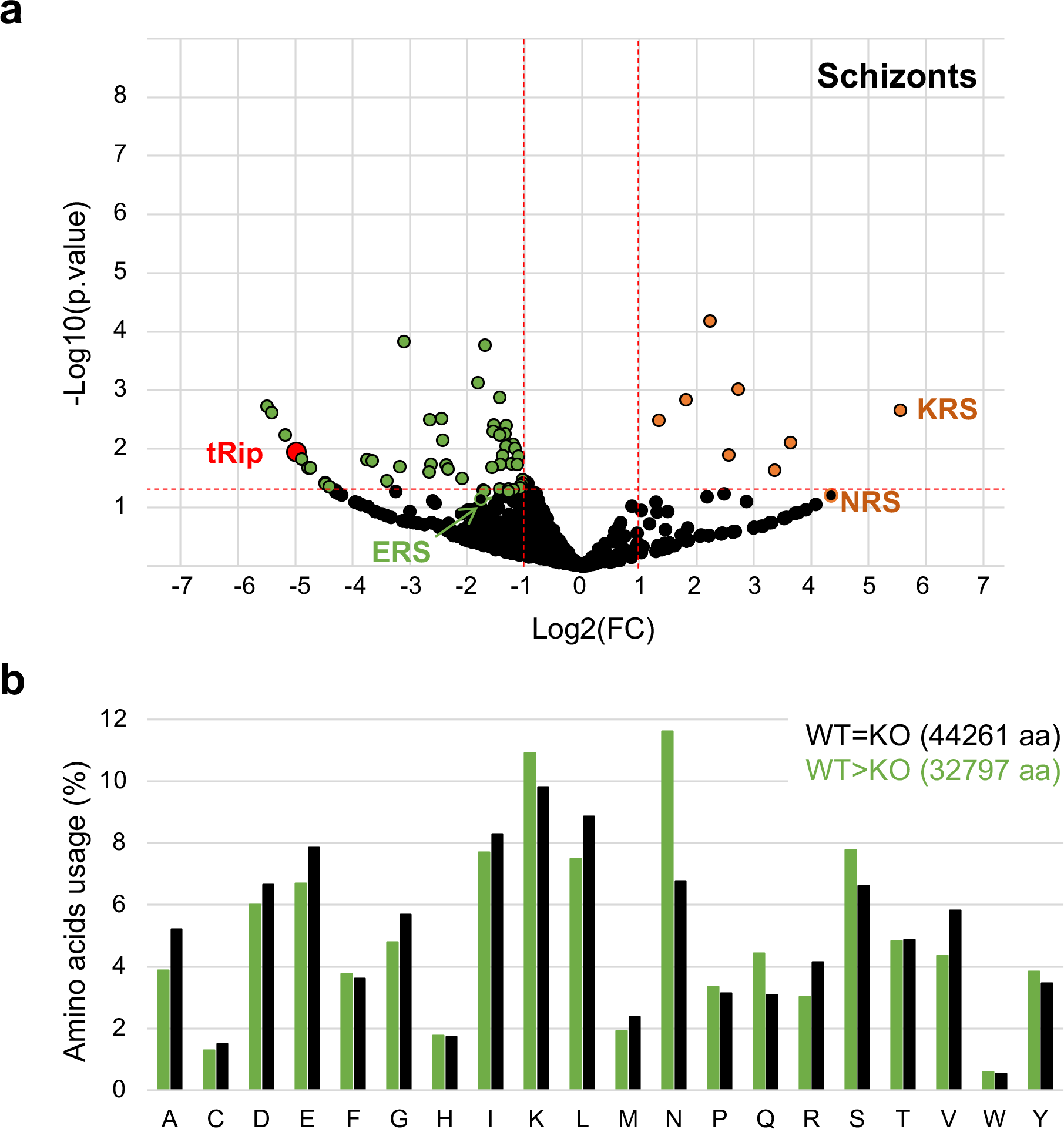
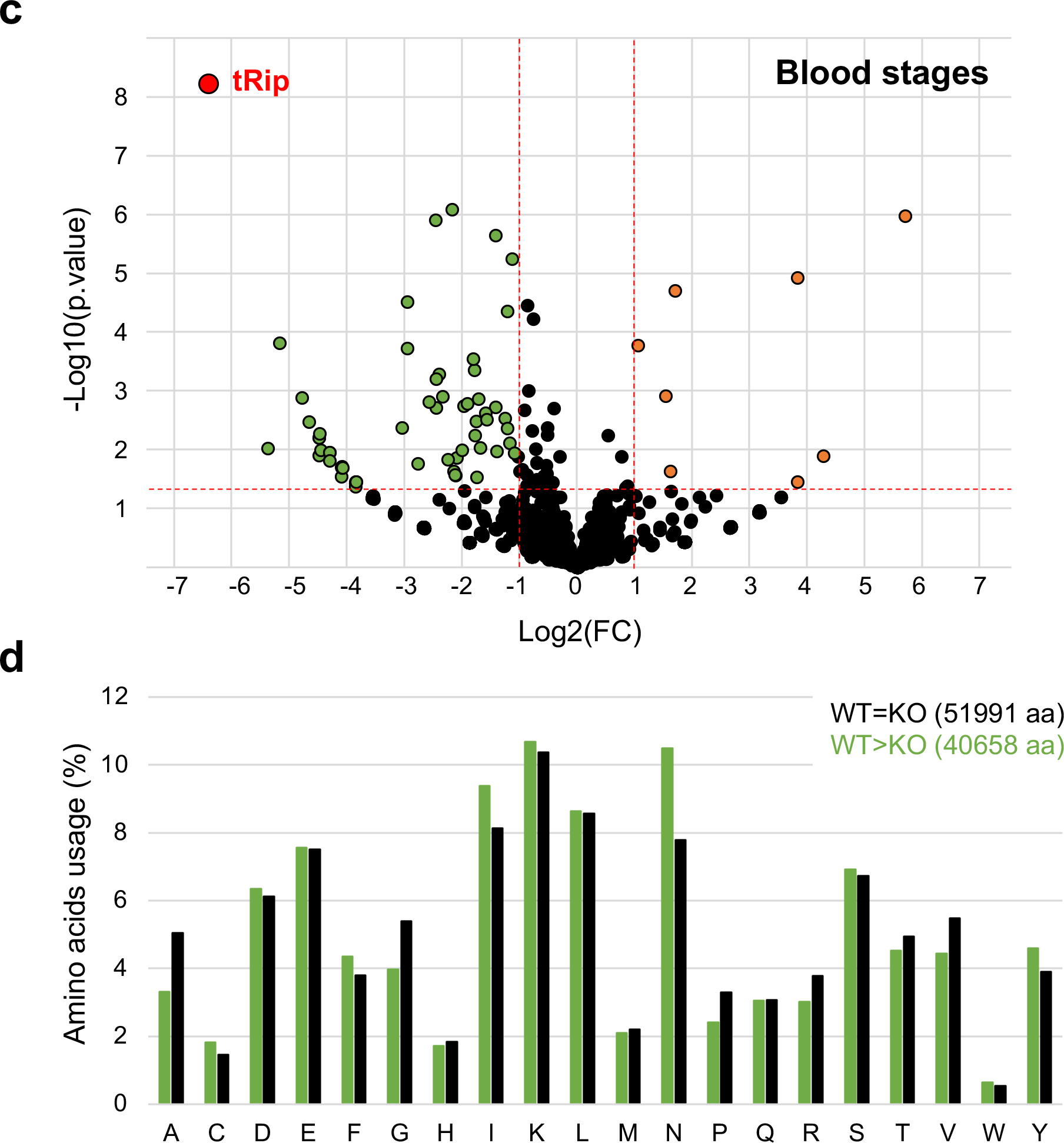
Comparative proteomic analysis and amino acid usage of proteins identified in WT and tRip-KO samples containing schizonts (a and b) or all blood stages (c and d). (**a**) Volcano plot of all quantified proteins from WT and tRip-KO parasites displaying the relationship between statistical significance (−log10(*p-value*), y-axis) and log fold change (FC) of each protein (log2(FC), x-axis). Statistics are based on three independent experiments (Supplementary Fig. S4). Deregulated proteins in tRip-KO parasite compared to the WT parasite are shown in orange (up-regulated, FC≥2 and *p-value* ≤ 0.05) and green (down-regulated, FC≤-1 and *p-value* ≤ 0.05). tRip is shown in red and black dots represent proteins with no significant change. ERS, KRS and NRS correspond to glutamyl-, lysyl- and asparaginyl-tRNA synthetases; ERS and NRS are shown as black circles surrounded by grenn and orange, respectively because their *p-values* are slightly greater than 0.05. (**b**) Comparison of amino acid usage (%) of proteins whose expression is stable in the tRip-KO parasite (the 100 most expressed proteins, black) and all proteins that are down-regulated in the tRip-KO parasite (green). Amino acids are designated by their one letter symbol and the total number of amino acids used in the analyses is specified. (**c**) Volcano plot of all quantified proteins from WT and tRip-KO parasites. Statistics are based on three independent experiments (Supplementary Fig. S6). (**d**) Comparison of amino acid usage (%) between proteins whose expression is stable in the tRip-KO parasite (the 150 most expressed proteins, black) and proteins down-regulated in the tRip-KO parasite (green).

### Amino acid compositions of tRip-KO down-regulated proteins in schizonts

Amino acid usage (percentage of amino acids) in the 100 most expressed stable proteins in the tRip-KO parasite was compared with that of the 49 significantly down-regulated proteins (Fig. 3b). There is no major increase in glutamate, methionine or glutamine usage in down-regulated proteins compared to the proteins efficiently expressed in the tRip-KO parasite, indicating that the dissociation of the two MSCs caused by tRip deletion does not directly affect protein synthesis in the mutant parasite. On the other hand, we clearly observe a 70% increase in asparagine in the proteins down-regulated in the tRip-KO parasite (Supplementary Fig. S5, 11.6% in downregulated proteins *versus* 6.8% stable proteins), thus, suggesting that the tRip-KO parasite has difficulty translating asparagine-rich proteins.

### Quantitative proteomics of tRip-KO *versus* WT in “blood stages” samples

Three biological replicates for tRip-KO and WT containing all blood stages (rings, trophozoites, and schizonts) were used for comparative proteomics (Supplementary Fig. S1b and Fig. S6). Triplicate proteomic data sets (KO and WT) were compared and identified between 617 and 1017 proteins each time. Sixty-nine proteins were significantly deregulated (at least a 2-fold change in abundance), 8 were up-regulated and 61 were down-regulated in the tRip-KO parasites compared to the WT (*p-value* < 0.05) (Fig. 3c). The few up-regulated proteins are especially rhoptry associated proteins (a specialized organelle in *Plasmodia*) and proteins involved in mobility and invasion (Supplementary Fig. S6), while down-regulated proteins were distributed across different functional families: Apart from the 20% of proteins with unknown function, most are involved in DNA replication (41%). Yet, none were common to the schizonts samples (compare Supplementary Tables S1 and S3). Amino acid usage in the tRip-KO down-regulated proteins shows also that these proteins contain more asparagine residues (10.5% at tRip-KO versus 7.8% at WT, i.e. 34% more) than the 150 most expressed stable proteins (Fig. 3d, and Supplementary Fig. S5), leading to the same conclusion as above: in the absence of tRip, asparagine-rich proteins are less efficiently translated.

The mRNAs encoding 3 down-regulated proteins (MCM7, ChAF1-C and one protein with unknown function) were quantified by qRT-PCR. They all showed a strong decrease in the tRip-KO parasite compared to the WT parasite (Supplementary Fig. S7b), suggesting that down-regulated proteins are controlled at the level of mRNA turnover.

### Identification of conserved asparagine-rich proteins in several *Plasmodia* strains

Since both “schizonts” and “blood stages” samples strongly suggest that the tRip-KO parasite is hindered in inserting asparagine into proteins, the complete proteomes of six *Plasmodium* lineages (*P. falciparum*-5476 proteins, *P. berghei*-5076 proteins, *P. yoelii*-6097 proteins, *P. chabaudi*-5222 proteins, *P. knowlesi*-5340 proteins and *P. vivax*-6708 proteins) as well as *Toxoplasma gondii* (control, 8322 proteins) were retrieved from ApiDB ^9^ and analyzed for their asparagine content (Supplementary Fig. S8a). *Plasmodium* proteins contain more asparagine residues than *Toxoplasma*. Yet, the *P. knowlesi* and *P. vivax* proteomes contain fewer asparagine residues (about 8 %) than the 4 other *Plasmodia* strains (about 12%) (Supplementary Fig. S8a and Fig. S5). Proteins were then ranked from highest to lowest asparagine contents to identify those that might be impacted by decreased asparagine decoding efficiency. For all proteomes, only the top 0.5% proteins were considered to search for conserved proteins with the highest asparagine content. Depending on the strain, 25 to 30 proteins were selected (Fig. 4). Besides many proteins with unknown functions and different AP2 domain transcription factors, only 2 proteins are strictly conserved in all strains. They correspond to the CCR4-associated factor-1 (CAF-1) and the poly-A binding protein-3 (PABP-3). Both share the same ligand (poly-A sequence) and a peculiar modular organization. They feature a high-complexity N-terminal domain and a long low-complexity C-terminal domain containing a very large amount of asparagine residues (Supplementary Figs. S8b and S8c). The proportion of asparagine varies locally between 35 and 60% in *P. falciparum, P. yoelii, P. chabaudi* and *P. berghei* and between 20 and 25% in *P. knowlesi* and *P. vivax* (i.e. at least 3 times higher than their respective average usage of asparagine). Yet these two proteins are not detected or by only a few spectral counts in our proteomic data (Supplementary Tables S1 and S3).

**Figure 4.**
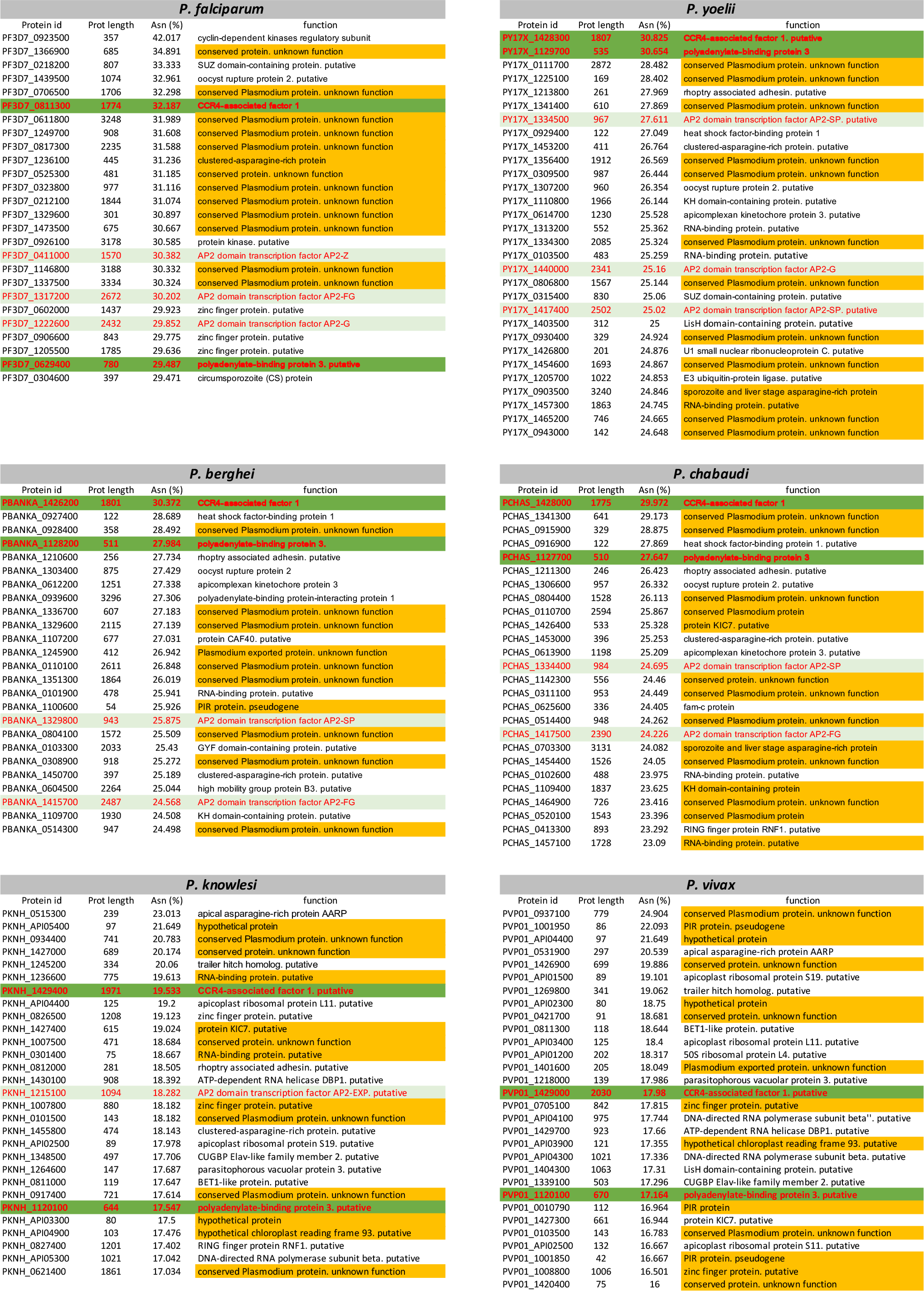
Top 0.5% of the most asparagine-rich proteins in 6 *Plasmodium* strains. Six *Plasmodium* species were selected: *P. falciparum, P. yoelii, P. berghei, P. chabaudi*, all featuring high asparagine usage (around 12%, Supplementary Fig. S8a) often localized in homorepetitions and *P. knowlesi* and *P. vivax* which show lower asparagine usage (around 8%, Supplementary Fig. S8a) and no homorepeats. Proteins highlighted in yellow are proteins of unknown function, light green indicate AP2 family regulatory proteins and dark green show the two proteins conserved in all 6 species, regardless of their asparagine usage.

## Discussion

To achieve efficient and specific protein synthesis, codon usage of mRNA must be balanced with the availability of corresponding aminoacylated tRNAs in the cell (i.e.^10–13^). Any discrepancy can affect the rate of protein elongation in ribosomes and result in pauses in translation that lead to mRNA degradation ^14^. It is interesting to note that blood contains reticulocytes, precursors of RBCs, which are very active in translation and therefore rich in tRNAs ^15^. Both reticulocytes (about 2%) and mature red blood cells (RBC) can be infected by *Plasmodia*. While reticulocytes are characterized by a dynamic protein synthesis, RBC are hyper-specialized anucleate cells that retain about 10 % of the protein synthesis observed in reticulocytes ^16^. Despite their scarcity, *P. berghei* ^17^, *P*.*chabaudi* ^18^, *P. yoelii* ^19^ and *P. vivax* ^20^ have been shown to preferentially invade these tRNA-rich reticulocytes compared to mature RBCs.

Moreover, by comparing the proteomes of wild-type and tRip-KO *P. berghei*, we observed in two independent experiments that down-regulated proteins in the tRip-KO parasite are asparagine-rich proteins. Asparagine is the most used amino acid in the *P. berghei* proteome and is often found in long homorepeats ^21,22^. This asparagine abundance is also found in *P. falciparum, P. chabaudi, P. yoelii* and to a lesser extent in *P*.*knowlesi* and *P. vivax* (Supplementary Fig. S8a), however proteins in the latter two parasites do not contain long asparagine repeats. Based on these results, we propose that import of host tRNA^Asn^ by tRip would ensure correct translation of asparagine-rich protein domains. In other words, the absence of tRNA import in the tRip-KO parasite would prevent the accumulation of host tRNA^Asn^ and would thus explain why asparagine-rich proteins are poorly translated. This hypothesis is strongly supported by several observations: (i) asparaginyl- and lysyl-tRNA synthetases are overexpressed in the proteomes of tRip-KO schizonts (Fig. 3a, Supplementary Fig. S4). Both aaRSs aminoacylate tRNAs that are most used to synthesize *P. berghei* proteins. In general, increased expression of aminoacyl-tRNA synthetase correlates with a decrease in cognate aminoacyl-tRNA and/or amino acid ^23,24^. This observation suggests, that tRNA^Asn^ and tRNA^Lys^ levels are low in tRip-KO schizonts. (ii) Furthermore, mammalian (human) tRNA^Asn^ is among the tRNAs with the highest affinity for *P. falciparum* tRip, along with tRNA^Ser^ _AGA_ and tRNA^Leu^ _hGA_ (Fig. 1) ^2^; It is reasonable to assume that the tRNAs with the highest affinity for tRip are also those that are most efficiently imported into the parasite. (iii) Finally, compensation for low tRNA^Asn^ by tRip-mediated tRNA import is only possible if the host tRNA^Asn^ is a substrate for the parasite AsnRS. This is indeed the case (Fig. 2) indicating that once in the parasite, at least some host tRNA^Asn^ isoacceptors can be efficiently aminoacylated and used in parasite mRNA translation.

Two asparagine-rich proteins stand out and are conserved in the six *Plasmodium* species analyzed (Fig. 4): the CCr4-associated factor 1 (CAF-1), involved in the poly-(A) decay of mRNAs ^25^, and to the poly-(A) binding protein-3 (PABP3). Not only are both proteins involved in poly-(A) recognition, but they also adopt a common global structure with long asparagine-rich C-terminal domains found only in *Plasmodia* (Supplementary Fig. S8b). While PABP3 has not yet been characterized, CAF-1 has been studied in detail in *P. falciparum*. CAF-1 is an essential nuclease that degrades mRNAs enriched with non-optimally decoded codons. It belongs to the eukaryotic CCr4-Not complex, which binds to the empty E-site of the ribosome and properly positions CAF-1 to initiate the decay of the 3′-poly(A)-tails of stalled mRNAs ^26^. However, the asparagine-rich C-terminal domain of *P. falciprum* CAF-1 is not essential and it has been shown that its deletion leads to the premature release of non-infectious merozoites *in vitro* ^27^ and to the inappropriate development of gametocytes *in vivo* ^28^. Interestingly, disruption of tRip and of the C-terminal domain of CAF-1 not only lead to the same phenotype: parasites have reduced infectivity and multiplication in the blood stage ^1,27^, but also results in overexpression of the same gene products involved in parasite egress and invasion: the rhoptry proteins, RAP1, RAP2/3 and the major surface proteins AMA-1 and MSP1 (Supplementary Fig. S6 and Fig. S8d). It suggests that tRip and CAF-1 may be linked in their cellular functions, i.e. in the tRip-KO parasite the absence of host tRNA^Asn^ import would hinder the synthesis of the asparagine-rich C-terminus of CAF-1, and lead to the specific and premature expression of mRNAs encoding the proteins responsible for merozoite release and infectivity.

The complex life cycle of *Plasmodium* is highly regulated, involving tight transcriptional and especially post-transcriptional controls each time the parasite moves from one stage to another ^29,30^. Translational regulation of specific proteins which depends on the availability of host tRNAs would enable efficient and rapid control of the transition from one stage to another during development. Failure to import host tRNAs could indicate to the parasite that the cell has run out of resources and that it’s time to leave and infect another cell. In this respect, tissue-specific variations in tRNA levels and in their post-transcriptional modifications ^31,32^ represent diverse sources of tRNAs for import. Depending on the available tRNAs isodecoders, their concentration and their post-transcriptional modifications in the host cell, tRNA import could differentially control parasite translation and enable it to develop optimally not only in blood, but also in liver or mosquito. In this model, host tRNA supply could play a major role in parasite development by modulating the translation efficiency of certain mRNAs, while complying with the “just-in-time” translation model ^29^.

## Methods

### Bioinformatics

Protein sequences as well as proteomes from all *Plasmodium* strains were retrieved from PlasmoDB. A home-made Python 2.7 script was used to calculate the asparagine content for each protein sequence. *Toxoplasma gondii* was the outgroup species.

### Parasite production

Four- to six-week-old female mice (C57BL/6), weighing approximately 20 g, were injected intraperitoneally with 200 µL of frozen infected red blood cells (10-15% parasitemia diluted in phosphate-buffered saline (PBS)) either with wild-type (WT, *Pb* gfpGOMO14) or with tRip-KO (*Pb* tRip-KO mCherry) malaria parasites derived from the *P. berghei* (ANKA strain). Parasitemia was monitored daily by cytometry (BD Accuri C6). Mice with parasitemia between 5 and 10% were selected. Therefore, in this study, only mice without or low symptoms were used. These parasitemia levels were reached 3-6 days after parasite injection. Mice were put to sleep and blood was collected by intracardiac puncture (about 1 to 1.5 mL) and parasites were directly purified (all blood stages) or cultured for 24 hours in RPMI at 37°C under 5% CO2 (only schizonts) (Supplementary Fig. S1b) as described in ^33^.

infected blood was filtered through a Plasmodipur filter (Europroxima) to remove mouse leukocytes, centrifuged for 10 min at 450 g and recovered in 4 mL of RPMI. A 7.2 mL cushion of 60% isotonic Percoll was gently pipetted under the red blood cells and the tube was centrifuged for 20 min at 1450 g (swinging buckets), to separate infected red blood cells (iRBC) at the Percoll/RPMI interface from non-infected RBC at the bottom of the tube. Parasitized RBCs were recovered in 2 tubes, washed 3 times in 1 mL of PBS and combined with 200 µL each of PBS. Infected RBCs were then lysed with 0.02% saponin for 5 min in ice (in 400 µL). Free parasites were recovered by centrifugation for 5 min at 2000 g and washed in 500 µL of PBS. The two pellets were resuspended either in 50 µL of protein loading buffer and stored at -80°C until mass spectrometry analysis, or as is and placed at - 80°C for RNA preparation. All experiments were performed in accordance with the ARRIVE guidelines and regulations under the project license for animal experimentation APAFIS#11124-2018010312571506 v2.

### Mass spectrometry (MS) and data analyses

Approximately 10 µg of protein was obtained from the half of the blood (~0.75 mL of a 5-10% infected mouse). Protein concentrations were determined by Bradford assay using bovine serum albumin as the standard. Proteins were precipitated, reduced, and alkylated as described in ^3^. After digestion overnight with sequencing grade porcine trypsin (300 ng, w/w, Promega, Fitchburg, MA, USA), the generated peptides were analyzed either using Easy-nanoLC-1000 system coupled to a Q-Exactive Plus mass spectrometer (Thermo Fisher Scientific, Germany) with 160-minutes gradients (blood stages) or a NanoLC-2DPlus system (nanoFlexChiP module; Eksigent, ABSciex, Concord, Ontario, Canada) coupled to a TripleTOF 5600 mass spectrometer (ABSciex) operating in positive mode with 120-minutes gradients (schizonts). Data were searched using the Mascot algorithm (version 2.6.2, Matrix Science) against the Uniprot database with *P*.*Berghei* taxonomy (release 2021_03, 4 927 sequences) with a decoy strategy. The resulting .dat Mascot files were then imported into Proline version 2.0 software ^34^ to align the identified proteins. Proteins were then validated with Mascot pretty rank equal to 1, and 1% false discovery rate (FDR) on both peptide spectrum matches (PSM) and protein sets (based on Mascot score).

For statistical analyses, raw Spectral Count values were imported into R (v. 3.5.0) where the number of spectra were first normalized using the DESeq2 median of ratio normalization method. A negative-binomial test using an edgeR GLM regression generated for each identified protein a p-value and a protein fold-change (FC). The R script used to process the dataset is published on Github ^35^. Proteins were statistically enriched or decreased with a p-value < 0.05 and a minimum fold change (FC) of 2 or 0.5, respectively. Mass spectrometry proteomic data will be deposited within the ProteomeXchange Consortium via the PRIDE partner repository ^36^ with the dataset ID PXD043916.

### RNA purification and QRT-PCR

Total RNA was extracted from parasites using the RNeasy Mini Kit (Qiagen) according to the manufacturer’s protocol and was subjected to DNAse treatment with the Rapid OUT DNA removal kit (Thermo Scientific). Total RNA was analyzed and quantified on Bioanalyser (puce PICO). Each sample was reverse transcribed in a 20 µL reaction volume containing 10 µL (0.16 to 0.5 μg) of RNA, using the SuperScript II reverse transcriptase (Invitrogen) according to the manufacturer’s protocol. The mRNAs levels were measured by RT-PCR (25 µL containing 4 µL of cDNA) on a CFx94 (Bio-Rad) using the Syber Green kit (Thermo Scientific). Oligonucleotides used for qRT-PCR are listed in the Supplementary Fig. S7a. qRT-PCR reactions were designed according MIQE guidelines ^37^, the specificity of the oligonucleotides was validated, and the amplification efficiencies of the primer sets are all between 90 and 110% and r^2^ values greater than 0.96. The mRNAs levels were calculated according to the ∆Cq method and normalized by the mRNA level of both the EF1-α and Hsp70 in each sample. Raw data are indicated as mean of three measurements, and results were expressed as the average of 3 biological samples ± standard error of the mean (SEM).

### Purification of aminoacyl-tRNA synthetases and aminoacylation assays

Recombinant *P. falciparum* and *H. sapiens* NRS and DRS were cloned, expressed and purified as described in references ^38,39^ respectively. The gene encoding the *P. falciparum* TyrRS was amplified by PCR from *P. falciparum* cDNA and cloned into pQE30 (Qiagen) and the plasmid encoding the *H. sapiens* YRS was a gift from P. Schimmel. Both *P. falciparum* and *H. sapiens* YRS were expressed and purified as described for NRS ^38^. Purified enzymes are shown in Supplementary Fig. S2c. *H. sapiens* crude tRNAs was prepared as described in ^2^.

Aminoacylation assays were performed under the same conditions: at 37 °C for 2, 4 and 6 min in 50 mM HEPES-KOH (pH 7.5), 20 mM KCl, 10 mM MgCl_2_, 2 mM ATP, 6 µM total tRNA from HeLa cells, in the presence of 20 µM of the corresponding L-[^14^C]-amino acid (Perkin Elmer). AaRSs were diluted in 100 mM HEPES-KOH pH 7.5, 1 mM DTT, 5 mg/mL BSA, and 10% glycerol and used at a final concentration of 200 nM in the aminoacylation assay. The aminoacylation plateaus are the mean of 3 independent experiments ± standard deviation (SD).

## Aknowledgments

We are grateful to Philippe Hammann, Lauriane Kuhn, and Béatrice Chane Woon Ming for LC-MS/MS analysis, Fabrice Auge and Eric Marois for animal experiments, and to Dr Alain Lescure and Prof Tamara Hendrickson for providing comments on this manuscript.

This work was performed under the framework of the Interdisciplinary Thematic Institute IMCBio, as part of the ITI 2021-2028 program of the University of Strasbourg, CNRS and Inserm. It was supported by IdEx Unistra (ANR-10-IDEX-0002), by SFRI-STRAT’US project (ANR 20-SFRI-0012), and EUR IMCBio (IMCBio ANR-17-EURE-0023) under the framework of the French Investments for the Future Program », by previous Labex NetRNA (ANR-10-LABX-0036), by the CNRS and the Université de Strasbourg, IdEx “Equipement mi-lourd” (2015) and Equipement d’Excellence (EquipEx) I2MC (ANR-11-EQPX-0022), and by the Fondation pour la Recherche Médicale (FRM) (grant number FDT201704337050) to Marta Cela.

## Authors Contributions

MP, DK and MC performed experiments, data acquisition, and analysis, JC performed mass spectrometry analysis and LD provided computer programming. MF managed the conception, design, interpretation of data and funding acquisition and wrote, reviewed, and edited the manuscript.

## Data availability statement

All relevant data are within the manuscript and its Supporting Information files.

## Additional information

### Competing Interest Statement

The authors declare no conflict of interest.

1 pdf with 5 supplementary Figures and 1 excel document containing 3 supplementary data sets named Supplementary Figs. S4, S5 and S6.

