## Supplemental Figures S1-S2-S3-S7-S8 for "Comparative proteomics uncovers low asparagine insertion in *Plasmodium* tRip-KO proteins"

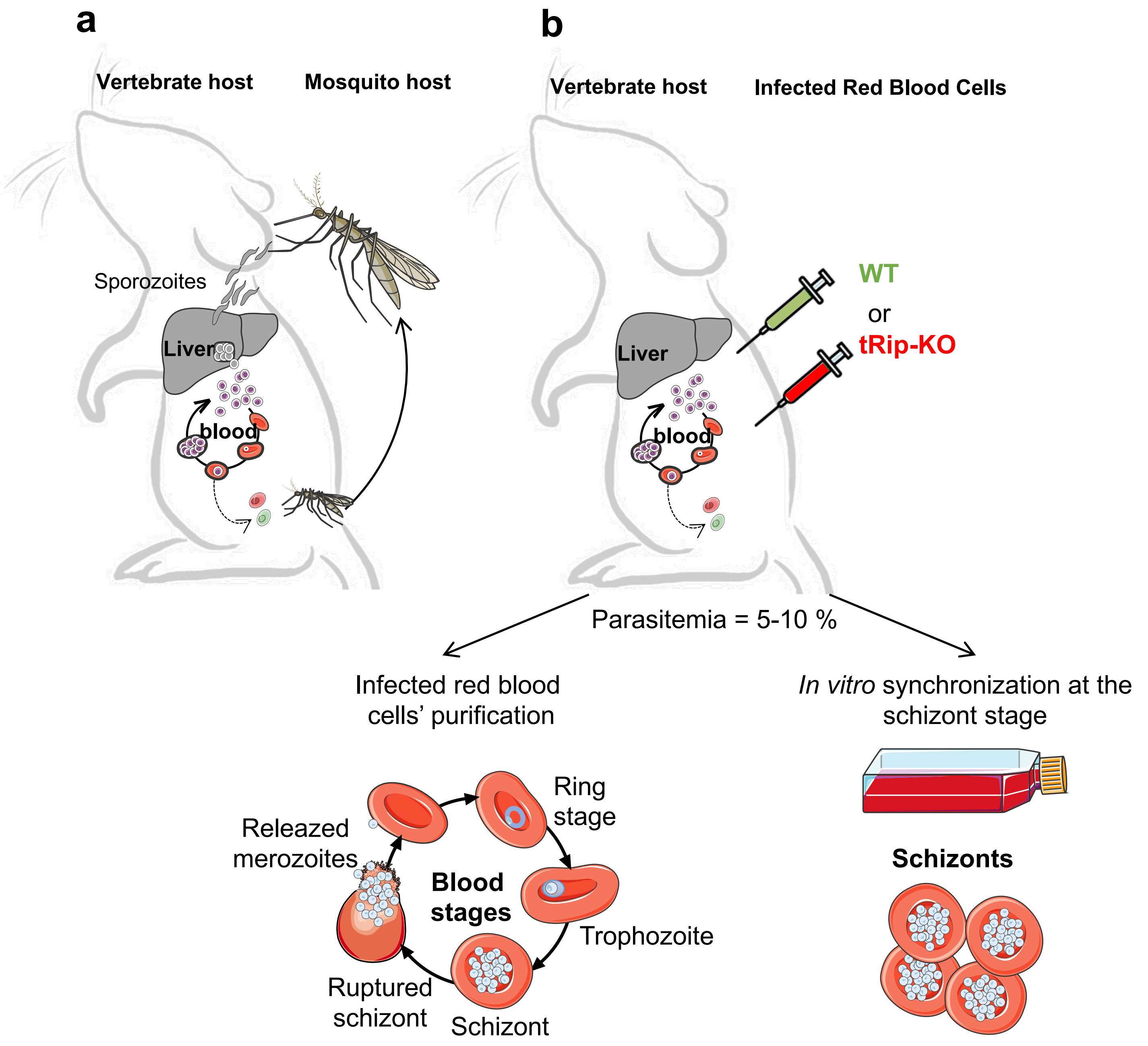


**Supplementary Fig. S1. Mice infections and sample preparation.** (**a**) *Plasmodium* life cycle. The cycle of vertebrate infection starts when a mosquito injects sporozoites into the host. Sporozoites invade hepatocytes (liver stage), multiply, and produce tens of thousands of merozoites per infected hepatocyte[1]. Merozoites exit the liver and develop in red blood cells (blood stages) to produce 10 to 30 new merozoites per intra-erythrocytic cycle[2]. Some of the erythrocytic merozoites differentiate into gametocytes (sexual forms, dashed lines)[3]. Fertilization takes place in mosquitoes where gametocytes are ingested during a blood meal. In 8-15 days, sporozoites invade the mosquito salivary glands, completing the cycle. (**b**) Alternatively, mice are infected with frozen stocks of red blood cells infected with wild-type (GFP) or tRip-KO (mCherry) *P. berghei* parasites. Three to six days later (5-10% parasitemia), infected blood is collected and used as is for analysis (all blood stages) or synchronized *in vitro* to the schizont stage[4]. "All blood stages" samples contain rings, trophozoites and merozoites.

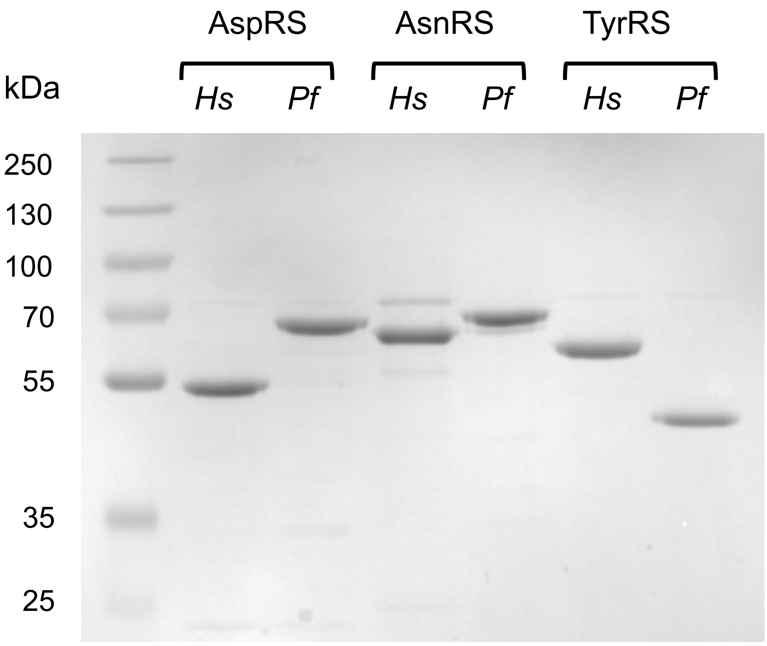


**Supplementary Fig. S2.** Gel analysis of aminoacyl-tRNA synthetases. One µg of each recombinant enzyme aspartyl- (DRS), asparaginyl- (NRS) and tyrosyl-(YRS) tRNA synthetases from *H. sapiens* (*Hs*) or *P. falciparum* (*Pf*) was run on a 10 % SDS PAGE.

**a**

| Genotype | Sample | Average Ct (triplicate) MCM7 | Average Ct (triplicate) Hsp70 | Average Ct (triplicate) EF1-α | ΔCt MCM7 | ΔCt Hsp70 | ΔCt EF1-α | Efficiency MCM7 | Efficiency Hsp70 | Efficiency EF1-α | Mean Hsp70/EF1-α | RQ MCM7 | **Mean RQ MCM7** | SD | **SEM** |
|---|---|---|---|---|---|---|---|---|---|---|---|---|---|---|---|
| WT | 2 | 19,82 | 19,77 | 17,86 | -1,10 | -0,59 | 0,88 | 2,14 | 1,50 | 0,54 | 0,90 | 2,37 | | | |
| WT | 3 | 20,73 | 21,54 | 16,21 | -0,19 | 1,18 | -0,77 | 1,14 | 0,44 | 1,71 | 0,87 | 1,32 | **1,77** | 0,49 | **0,28** |
| WT | 4 | 20,79 | 20,96 | 17,50 | -0,13 | 0,60 | 0,52 | 1,09 | 0,66 | 0,70 | 0,68 | 1,62 | | | |
| KO | 1 | 21,87 | 19,99 | 16,73 | 0,95 | -0,37 | -0,25 | 0,52 | 1,29 | 1,19 | 1,24 | 0,42 | | | |
| KO | 2 | 21,27 | 19,75 | 16,30 | 0,35 | -0,61 | -0,68 | 0,78 | 1,52 | 1,60 | 1,56 | 0,50 | **0,62** | 0,25 | **0,14** |
| KO | 4 | 21,04 | 20,14 | 17,27 | 0,12 | -0,22 | 0,29 | 0,92 | 1,16 | 0,82 | 0,97 | 0,94 | | | |
| Mean | | 20,92 | 20,36 | 16,98 | | | | | | | | | **p-value = 0,03** | | |

| Genotype | Sample | Average Ct (triplicate) ChAF1C | Average Ct (triplicate) Hsp70 | Average Ct (triplicate) EF1-α | ΔCt ChAF1C | ΔCt Hsp70 | ΔCt EF1-α | Efficiency ChAF1C | Efficiency Hsp70 | Efficiency EF1-α | Mean Hsp70/EF1-α | RQ ChAF1C | **Mean RQ ChAF1C** | SD | **SEM** |
|---|---|---|---|---|---|---|---|---|---|---|---|---|---|---|---|
| WT | 2 | 17,89 | 19,37 | 15,75 | -3,29 | -2,68 | -2,32 | 9,75 | 6,42 | 4,98 | 5,65 | 1,72 | | | |
| WT | 3 | 24,23 | 26,49 | 23,51 | 3,06 | 4,44 | 5,45 | 0,12 | 0,05 | 0,02 | 0,03 | 3,70 | **2,70** | 0,88 | **0,51** |
| WT | 4 | 21,34 | 23,53 | 19,75 | 0,16 | 1,48 | 1,69 | 0,89 | 0,36 | 0,31 | 0,33 | 2,67 | | | |
| KO | 1 | 23,80 | 23,57 | 16,11 | 2,63 | 1,52 | -1,96 | 0,16 | 0,35 | 3,88 | 1,16 | 0,14 | | | |
| KO | 2 | 19,23 | 18,95 | 15,72 | -1,95 | -3,10 | -2,35 | 3,85 | 8,59 | 5,08 | 6,61 | 0,58 | **0,48** | 0,27 | **0,15** |
| KO | 4 | 20,56 | 20,41 | 17,55 | -0,62 | -1,64 | -0,52 | 1,53 | 3,12 | 1,43 | 2,11 | 0,72 | | | |
| Mean | | 21,18 | 22,05 | 18,07 | | | | | | | | | **p-value = 0,02** | | |

| Genotype | Sample | Average Ct (triplicate) UF | Average Ct (triplicate) Hsp70 | Average Ct (triplicate) EF1-α | ΔCt UF | ΔCt Hsp70 | ΔCt EF1-α | Efficiency UF | Efficiency Hsp70 | Efficiency EF1-α | Mean Hsp70/EF1-α | RQ UF | **Mean RQ UF** | SD | **SEM** |
|---|---|---|---|---|---|---|---|---|---|---|---|---|---|---|---|
| WT | 2 | 19,91 | 19,78 | 16,19 | -2,09 | -0,61 | -0,72 | 4,26 | 1,53 | 1,64 | 1,59 | 2,68 | | | |
| WT | 3 | 21,13 | 21,74 | 17,98 | -0,87 | 1,35 | 1,07 | 1,83 | 0,39 | 0,48 | 0,43 | 4,23 | **3,23** | 0,77 | **0,44** |
| WT | 4 | 20,98 | 20,92 | 17,30 | -1,02 | 0,53 | 0,39 | 2,03 | 0,69 | 0,76 | 0,73 | 2,79 | | | |
| KO | 1 | 24,04 | 19,92 | 16,70 | 2,04 | -0,47 | -0,21 | 0,24 | 1,39 | 1,15 | 1,27 | 0,19 | | | |
| KO | 2 | 22,68 | 19,47 | 16,01 | 0,68 | -0,93 | -0,90 | 0,62 | 1,90 | 1,86 | 1,88 | 0,33 | **0,34** | 0,14 | **0,08** |
| KO | 4 | 23,26 | 20,54 | 17,26 | 1,26 | 0,15 | 0,35 | 0,42 | 0,90 | 0,78 | 0,84 | 0,50 | | | |
| Mean | | 22,00 | 20,40 | 16,91 | | | | | | | | | **p-value = 0,0046** | | |

| **Gene ID and name** (expression status in tRip-KO) | **Primers** | Primer efficiency | $r^2$ |
|---|---|---|---|
| **PBANKA_0803100** DNA replication licensing factor MCM7 (down regulated) | Forward CAACCATTTCCCCTGAACTA<br>Reverse CACTATCTCTTACCCTTGCC<br>this study | 105.4 % | 0.965 |
| **PBANKA_0203000** chromatin assembly 1 (down regulated) | Forward GGACATTTTGCATCTGGAAG<br>Reverse GCATCTAACTCTTCACCGAT<br>this study | 99.8 % | 0,98 |
| **PBANKA_1029400** unknown function (down regulated) | Forward TAGCTACTTGTTTACCGCAA<br>Reverse TGAGGCCCCTTTAATTACTCTGT<br>this study | 96.0% | 0.990 |
| **Reference genes** | | | |
| **PBANKA_1133300** EF1-α | Forward TGGAACCACCCAAAAGACCA<br>Reverse ACAACAGCAGATGGAGCGAA<br>Tokunaga et al., 2019 | 100.7% | 1 |
| **PBANKA_0914400** Hsp70 | Forward AGAGAAGCAGCTGAAACAGC<br>Reverse TCCCTTTAATAAATCATGGC<br>Sanyal et al., 2013 | 106.9% | 0.999 |

**RNA analyses**

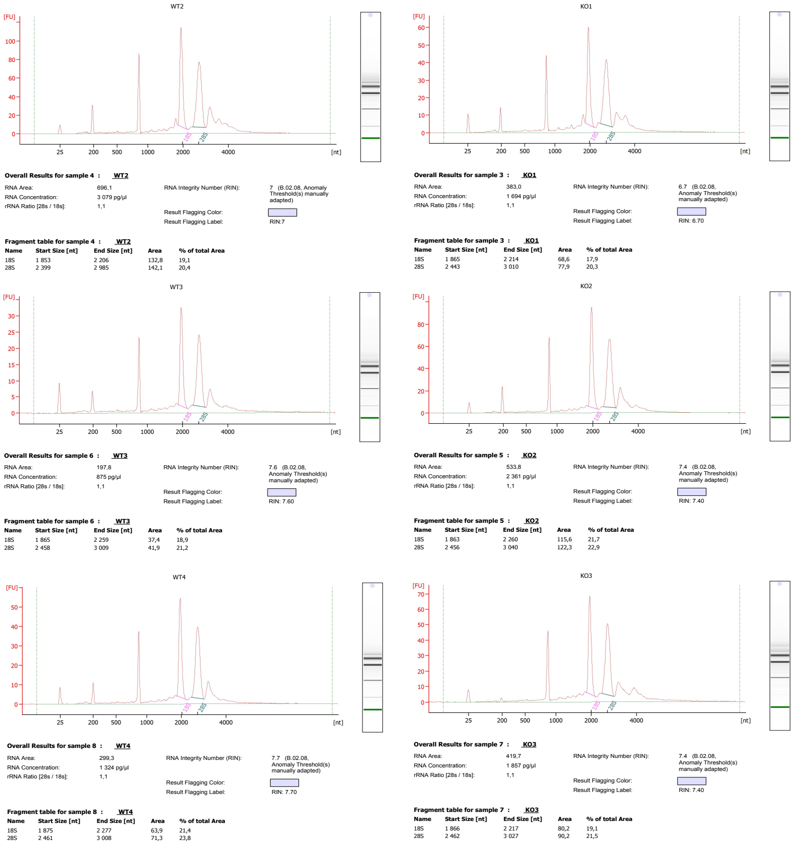


**b**

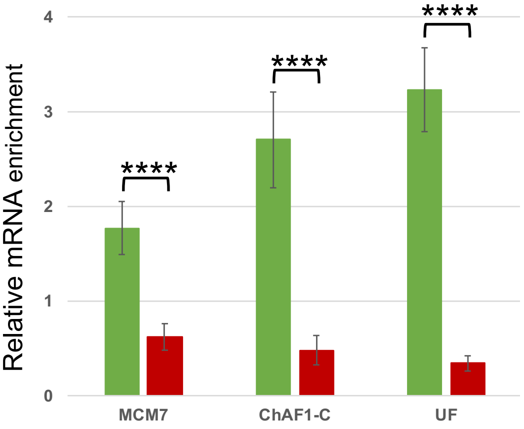


**Supplementary Fig. S7. qRT-PCR analysis of the mRNAs coding for deregulated proteins in the tRip-KO parasite.** The relative enrichment of mRNA was measured by qRT-PCR and determined by the $\Delta\Delta Ct$ method using both EF1-a (PBANKA_1133300) and Hsp70 (PBANKA_0914400) as normalizers. mRNAs are coding for down-regulated proteins: the DNA replication licensing factor MCM7 (MCM7, FC = 5.7, *p-value* =$1.3 \times 10^{-6}$), chromatin assembly factor 1-subunit C (ChAF1-C, FC = 2.2, *p-value* = $5.7 \times 10^{-6}$) and a protein with unknown function (UF, FC = 3.6, *p-value* = $2.9 \times 10^{-4}$). (**a**) QRT-PCR raw data. For each "all blood stages" samples, qRT-PCR data the primers as well as the quality control of the RNA samples are shown. (**b**) Error bars represent the standard errors of the means (SEM) of three independent experiments performed on "All blood stages" samples, each corresponding to triplicate qRT-PCR measurements (S4 Table). Asterisks indicate statistically significant differences with the WT control mRNA: **** *p-value* < 0.0001 based on Student's *t* test.

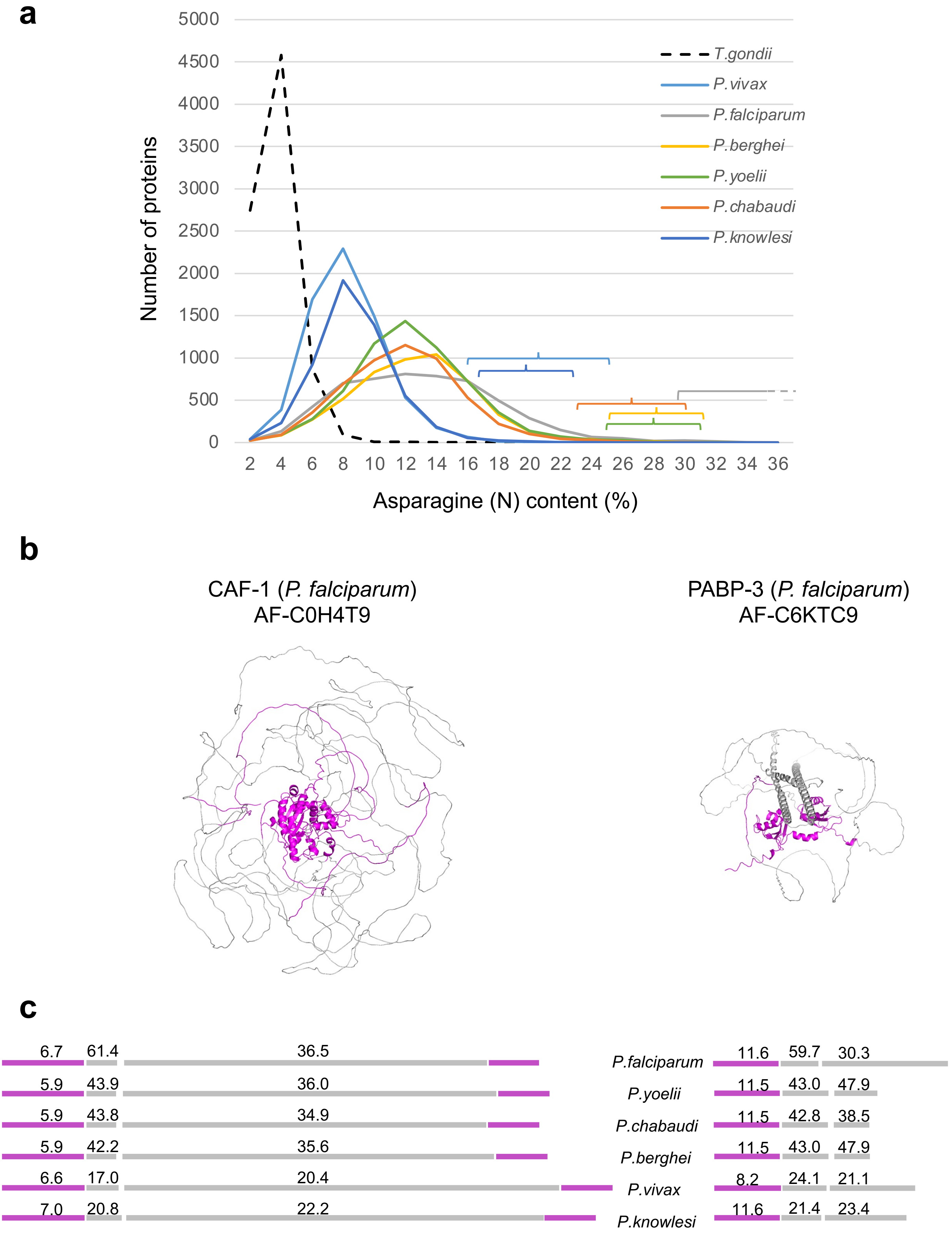

a
5000
4500
4000
3500
3000
2500
2000
1500
1000
500
0
Number of proteins
T.gondii
P.vivax
P.falciparum
P.berghei
P.yoelii
P.chabaudi
P.knowlesi
2 4 6 8 10 12 14 16 18 20 22 24 26 28 30 32 34 36
Asparagine (N) content (%)
b
CAF-1 (P. falciparum)
AF-C0H4T9
PABP-3 (P. falciparum)
AF-C6KTC9
c
6.7 61.4 36.5
5.9 43.9 36.0
5.9 43.8 34.9
5.9 42.2 35.6
6.6 17.0 20.4
7.0 20.8 22.2
P.falciparum
P.yoelii
P.chabaudi
P.berghei
P.vivax
P.knowlesi
11.6 59.7 30.3
11.5 43.0 47.9
11.5 42.8 38.5
11.5 43.0 47.9
8.2 24.1 21.1
11.6 21.4 23.4

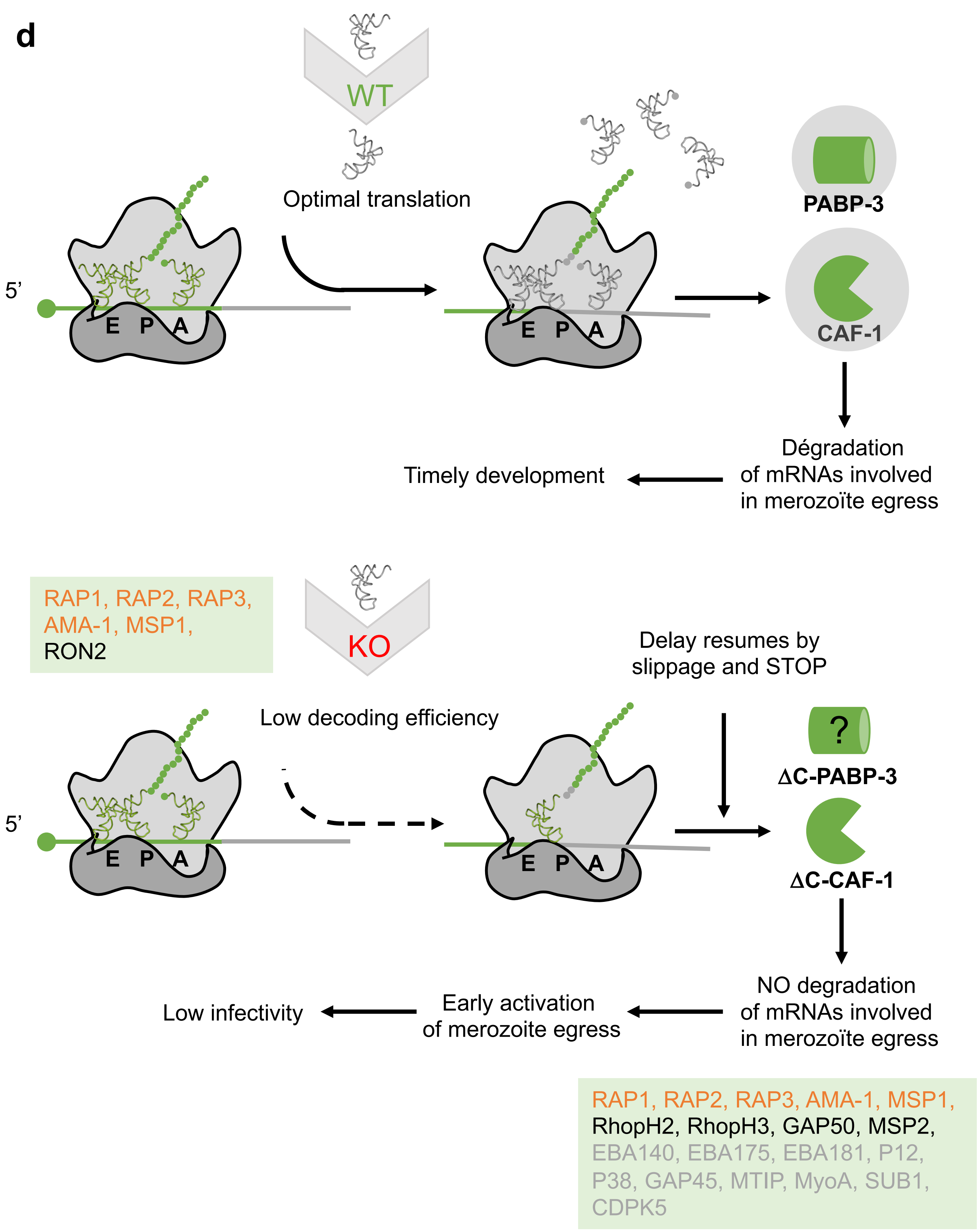
d
WT
Optimal translation
5'
E P A
E P A
PABP-3
CAF-1
Dégradation
of mRNAs involved
in merozoïte egress
Timely development
RAP1, RAP2, RAP3,
AMA-1, MSP1,
RON2
KO
Low decoding efficiency
Delay resumes by
slippage and STOP
5'
E P A
E P A
?
ΔC-PABP-3
ΔC-CAF-1
NO degradation
of mRNAs involved
in merozoïte egress
Early activation
of merozoite egress
Low infectivity
RAP1, RAP2, RAP3, AMA-1, MSP1,
RhopH2, RhopH3, GAP50, MSP2,
EBA140, EBA175, EBA181, P12,
P38, GAP45, MTIP, MyoA, SUB1,
CDPK5

**Supplementary Fig. S8. Asparagine frequency in *Plasmodium* proteins and functional link between tRip and CAF-1.** (**a**) Proteomes of *P. falciparum, P. yoelii, P. berghei, P. chabaudi*, *P. knowlesi* and *P. vivax* were compared and the proteome of *Toxoplasma gondii* was used as a negative control[5]. For each species, the number of proteins is indicated for a given asparagine content. Brackets show the top 0.5 % proteins containing the highest percentage of asparagine residues. (**b**) AlphaFold models of *P. falciparum* CAF-1 and PABP-3 (10). Both models show conserved domains (pink) and an unfolded asparagine-rich *Plasmodium* specific C-terminal extensions (grey). (**c**) Domain alignments of CAF-1 and PABP-3. The corresponding domains are shown with the same color code and the corresponding percentage of asparagine. (**d**) In WT parasites, the asparagine-rich C-terminal domains of *Plasmodium* CAF-1 and PABP-3 are efficiently translated thanks to $tRNA^{Asn}$ provided by tRip-mediated tRNA import from the host. Full-length CAF-1 can thus down-regulate specific mRNAs including those involved in egress and invasion that adequately control parasite development. On the contrary, in the tRip-KO parasite, translation of the C-terminal domains of CAF-1 and PABP-3 is hindered by the absence of host $tRNA^{Asn}$. The delay in translation can lead to frameshifting and the occurrence of stop codons. The resulting DC-CAF-1 protein can no longer regulate the translation of genes involved in parasite egress/invasion leading to early release of infectivity-deficient immature merozoites. Genes up-regulated in both mutants are highlighted in orange, they correspond to roptry-associated proteins 1, 2 and 3 (RAP1, PF3D7_1410400/PBANKA_1032100; RAP2/3, PF3D7_0501600/0501500, PBANKA_1101400), apical membrane antigen (AMA1, PF3D7_1133400/PBANKA_0915000) and merozoite surface protein 1 (MSP1, PF3D7_0930300/PBANKA_0831000). Up-regulated genes involved in merozoite's egress either in *P. falciparum* DC-CAF1 mutant or *P. berghei* tRip-KO parasite are listed inside green boxes: genes not in common between both strains are indicated in black: merozoite surface protein 2 (MSP2, PF3D7_0206800), glidosome associated protein 50 (GAP50, PF3D7_0918000/PBANKA_0819000), high molecular weight rhoptry proteins 2 and 3 (RhopH2, PF3D7_0929400/PBANKA_0830200 and RhopH3, PF3D7_0905400/PBANKA0416000) and Rhoptry neck protein 2 (RON2, PBANKA_1315700); Ten genes were up-regulated in the DC-CAF-1 mutant but were not detectable in blood stages proteomes. They correspond to: erythrocyte binding antigens 140, 175 and 181 (EBA140, PF3D7_1301600; EBA175, PF3D7_0731500; EBA181, PF3D7_0102500), 6-cysteine proteins P12 and P38 (PF3D7_0612700 and PF3D7_0508000), glideosome-associated protein 45 (GAP45, PF3D7_1222700), myosin A-tail interaction protein (MTIP, PF3D7_1246400/PBANKA_1459500), myosin A (MyoA, PF3D7_1342600), subtilisin-like protease 1 (SUB1, PF3D7_0507500) and calcium-dependent protein kinase 5 (CDPK5, PF3D7_1337800).
